# Extracellular release of a disintegrin and metalloproteinases orchestrates periodontal disease severity

**DOI:** 10.1101/2023.07.21.550016

**Authors:** Ahmad Aljohmani, Hakon Heinze, Federico Guillermo Gharzia, Bashar Reda, Ahmed Mohamed Mostafa Abdrabou, Sören Becker, Markus Bischoff, Matthias Hannig, Daniela Yildiz

**Affiliations:** Institute of Experimental and Clinical Pharmacology and Toxicology, PZMS, ZHMB, Saarland University; Clinic of Operative Dentistry, Periodontology and Preventive Dentistry, University Hospital Saarland, Saarland University; Institute of Medical Microbiology and Hygiene, Saarland University, Homburg, Germany; Medical Microbiology and Immunology Department, Faculty of Medicine, Mansoura University, Mansoura, Egypt

**Keywords:** proteolysis, metalloproteinase, exosomes, periodontal disease, matrix degradation

## Abstract

Periodontal diseases are amongst the most common pathologies worldwide with a high risk for the development of systemic complications. Periodontal disease is driven by oral pathogens such as *Porphyromonas gingivalis* and the release of inflammatory cytokines. These cytokines (e.g. TNF) or their receptors (IL-1R) are substrates of a disintegrin and metalloproteinases (ADAMs). In a comparative approach, we observed an increase of ADAM8 protein expression and activity in the sulcus fluid of periodontal disease patients correlating with the disease stage. In contrast, the induced ADAM10 expression was decreased. *In vitro* mechanistic studies revealed that both *Porphyromonas gingivalis* infection and the resulting cytokine release orchestrated the release of soluble ADAM8 by keratinocytes and neutrophils as soluble ectodomain and on exosomes, respectively. Furthermore, ADAM8 regulated the release of ADAM10 and MMP9, thereby potentially influencing wound healing and tissue destruction. Thus, the dysregulation of the cell-associated and extracellular ADAM proteolytic activity mainly driven by ADAM8 may be an essential regulatory element in periodontal disease onset and progression. This potential as novel local treatment option should be addressed in future translational studies.

## Introduction

Periodontal diseases are amongst the most common pathologies worldwide with an estimated global prevalence of 19 % and more than 1 billion cases worldwide (1). Chronic periodontitis may develop into systemic conditions based on hyper-inflammation and a disruption of the immune response, with the risk of endomyocarditis or changes of the gut microbiome (2). Periodontitis is initiated and driven by a dysbiosis of the oral microbiome resulting in an overactivation of the host immune response and changes in the cytokine network (3). Poor oral hygiene and lack of frequent biofilm disruption lead to the accumulation of quorum-sensing bacteria, such as *Fusobacterium nucleatum,* eliciting gingival inflammation and the proliferation of *Porphyromonas gingivalis* (*P. gingivalis*) (4). These oral bacteria manipulate the innate and adaptive immune response targeting especially neutrophils’ antimicrobial mechanisms to evade killing and to promote inflammation (5). The excess of reactive oxygen species, released matrix metalloproteinases (MMPs), and cytokines finally leads to collateral periodontal tissue loss and activation of osteoclasts resulting in bone loss (3, 4). Cytokines central to periodontitis are IL- 1β, IL-6, and TNF-α released by periodontal cells and innate immune cells, which again elicit the release of cytokines from T cells such as IFN-γ (3). These cytokines (TNF-α) or their corresponding receptors (IL-1R, IL-6R) are either substrates of a disintegrin and metalloproteinases (ADAMs) or induce their expression or activation (for review see (6)). ADAMs are multidomain transmembrane proteins, which are activated upon cleavage of the inhibitory pro-domain. Besides their proteolytic activity, ADAMs fulfill several non-proteolytic functions such as the interaction and regulation with integrins or with the extracellular matrix. In infectious diseases, ADAMs are involved in each step of the infection process, starting with the initial pathogen recognition and entrance, phagocytosis, and leukocyte recruitment up to regeneration (for review see (7)). ADAM proteases may not only act on the expressing cells themselves, but may be also released as soluble variants or on exosomes performing proteolytic cleavage in the extracellular matrix and at distinct cellular sites (8, 9). Earlier work demonstrated that ADAM8 is up-regulated in the saliva of periodontitis patients displaying proteolytic activity (10, 11), and its expression is enhanced by stimulation with *Fusobacterium nucleatum* in epithelial cells (12). Only a few studies have addressed the relevance of ADAM10 and ADAM17 in periodontal disease so far. For ADAM17, an up-regulation of the mRNA expression in gingival tissue was observed (13), and an increase of receptor activator of nuclear factor-κB ligand (RANKL) expression in osteoblast was linked to ADAM17 protein levels (14). Furthermore, inhibition experiments revealed that ADAM10 and ADAM17 may contribute to the release of chemokines from human gingival fibroblasts, thereby controlling the migration of leukocytes into the inflamed tissue (15). However, a comparative and mechanistic study of the ADAM-based proteolytic activity and their functional impact is missing so far.

We hypothesized that proteolytic activity of ADAMs released to the extracellular space could play an essential role in periodontal disease initiation and progression. In the recent study, we observed an increase in ADAM8 protein expression and proteolytic activity in the sulcus fluid of periodontal disease patients correlating with MMP9 release and the disease stage. Mechanistic *in vitro* studies revealed that the release of soluble ADAM8 by keratinocytes and exosomal ADAM8 by neutrophils are orchestrated by *P. gingivalis* and inflammatory cytokines. Furthermore, ADAM8 inhibition led to reduced MMP9 and ADAM10 release, pointing towards the regulation of tissue destruction. Thus, ADAM8 and its interaction partners could be novel treatment options for the prevention of periodontal disease initiation and progression, which should be addressed in future translational *in vivo* studies.

## Results

### The presence of ADAM8 and its proteolytic activity in gingival fluid increases with disease stage

To obtain first indications for a potential contribution of ADAMs to the progression of periodontal disease, protein expression of ADAM8, 10 and 17 was investigated in the sulcus fluid in healthy and periodontal disease patients. We observed a stage-dependent increase of ADAM8 total protein expression, shown by an increase of both the pro-from (120 kDa, only seen in 72 % patients) and mature form (90 kDa), and an increase of maturation (Fig. 1A-C, Suppl. Fig1A). The remnant form remaining after autocatalytic release of the extracellular domain was slightly but not significantly increased in all disease stages (Suppl. 1B). Interestingly, protein expression of ADAM10 as substrate of ADAM8 was strongly increased upon disease onset (stage I-II) but reduced at later stages of disease (Fig. 1D-E, Suppl. Fig1C, pro-from 100 kDa, mature form 70 kDa). This was further reflected on the maturation level (Fig. 1F), measured as ratio of the mature form to the pro-form. In contrast, ADAM17 did not show any change (Suppl. Fig. 1D-G, pro-form 130 kDa, mature form 100 kDa). It is important to note that healthy patients did not show any presence of these forms of ADAM8, 10 or ADAM17. As maturation does not directly indicate activation, we analyzed the activity of ADAM8 using a specific FRET-substrate including the cleavage site of CD23. While at early onset of disease (stage I-II) no changes in comparison to healthy patients were observed, activity massively increased in highly diseased patients (Fig. 1G, H).

**Figure 1.**
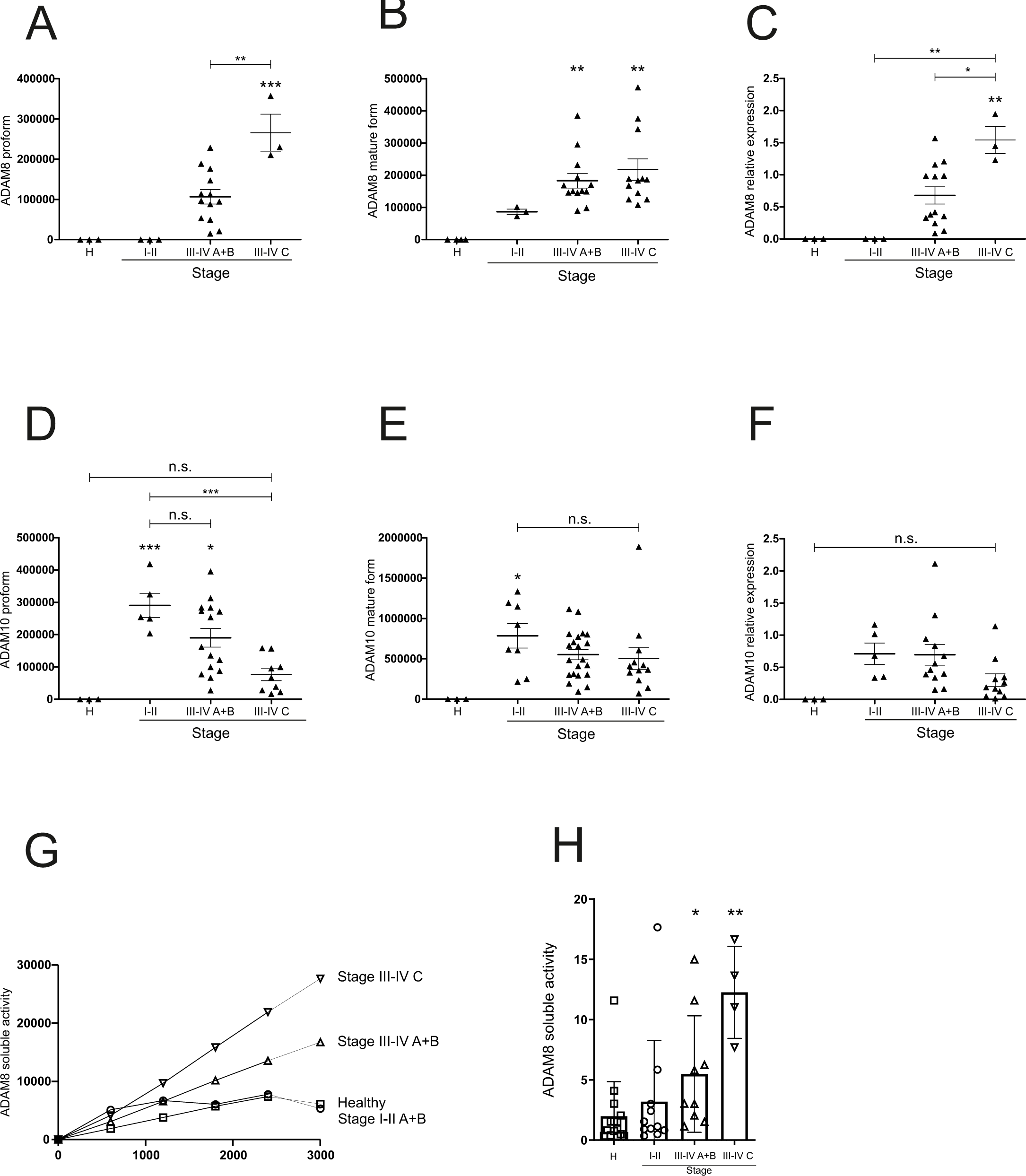
ADAM8 proteolytic activity in gingival fluid is associated with disease severity. (A-F) Sulcus fluid from patients with different disease stages were investigated by Western blot to analyze the expression and maturation of ADAM8 and ADAM10 using antibodies against the C-terminal domains. Band intensity was quantified by densitometry. (G-H) *S*ulcus fluid was resuspended in activity buffer and subjected to ADAM8 activity measurement using a FRET-based substrate (n=3 -12 per group). (A- F, H) Data are shown as mean ± SD, and activity over time in (G). Statistical analysis was performed with one-way ANOVA followed by Tukey post hoc test. Asterisks without line represent statistically significant differences to healthy patients, asterisks with line between groups (* p < 0.05, ** p < 0.01, *** p < 0.001).

### Porphyromonas gingivalis induces ADAM8 gene expression and release of ADAM8 to the extracellular matrix

Development of periodontitis is driven by an initial dysbiosis at the local site causing gingivitis, with *P. gingivalis* as central player of the red complex as a group of bacterial species associated with the clinical parameters of periodontitis. Therefore, we analyzed the expression of ADAM8 and ADAM10 in human primary keratinocytes upon infection with *P. gingivalis*. We observed an increase of ADAM8 pro-form and mature form, while the pro-form of ADAM10 was decreased (Fig. 2A-D, Suppl. Fig. 2A,B). As GAPDH could be only detected in the sulcus fluid of a few patients, we hypothesize that the observed expression and activity of ADAMs in patient samples may be mostly derived from cell-free structures. As mentioned before, ADAM8 is released as soluble ectodomain. Indeed, we observed an increase of the remaining ADAM8 remnant form upon infection (Fig. 2E). A second source of external activity could be the release of ADAM8-carrying exosomes. However, in the observed time frame only higher amounts of the remnant form were released on exosomes, but not of the mature form (Fig. 2F). Nevertheless, infection with *P. gingivalis* induced the release of mature ADAM10 on exosomes (Fig. 2G). Furthermore, FRET-based activity measurements revealed a strong increase of ADAM8 activity which was significantly inhibited by BK-1361 (16) (Fig. 2H). Further, we observed an increase in ADAM10 activity which was inhibited by GI254023X, but only slightly affected BK-1361 (Fig. 2I). Thus, the release of ADAM8 by oral keratinocytes as soluble ectodomain may contribute to the proteolytic pool observed in patient samples, while membrane-associated forms may be derived from different cellular sources.

**Figure 2.**
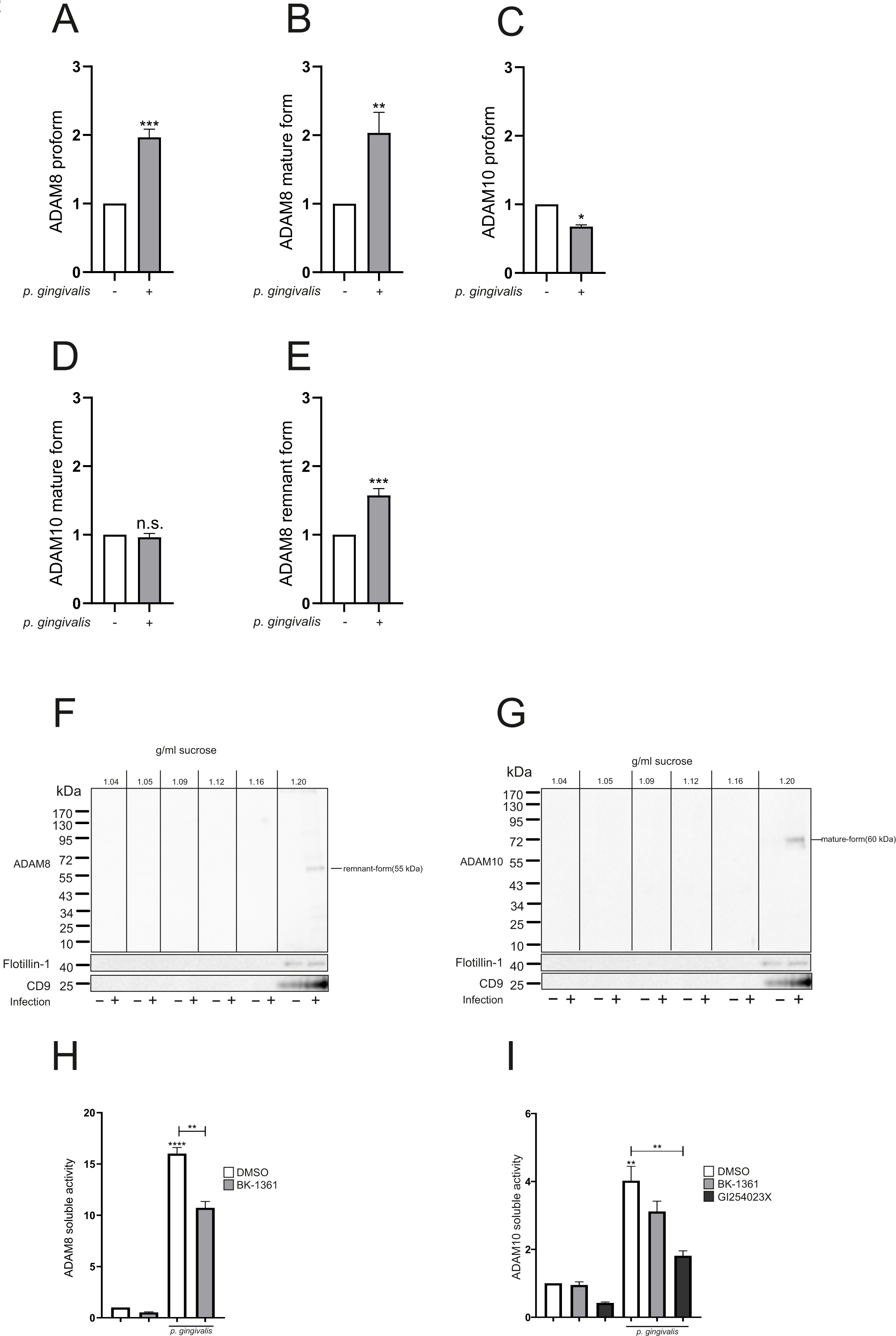
*P. gingivalis* induces ADAM8 expression and activity in human oral keratinocytes. Human primary keratinocytes (K2 cells) were grown until confluency and infected with *P. gingivalis* (2 h, 0.1 McF). (A-E) The cells were lysed, and the samples were analyzed by Western blot using antibodies against the C-terminal domains of ADAM8 and ADAM10, respectively. GAPDH served as loading control. Band intensity was quantified by densitometry and normalized to uninfected cells. (F,G) The medium of the infected and non-infected cells was subjected to differential centrifugation and density fractionation for exosome purification. The resulting pellets of different density were analyzed by Western blot for the expression of ADAM8 and ADAM10 as well as Flottilin-1 and CD9 as exosomal markers. Representative blots of three independent experiments are shown (H,I) K2 cells were pre- incubated with 10 µM BK-1361, 10 µM GI254023X or 0.01% DMSO (vehicle control) for 30 min and infected with *P. gingivalis* (2 h, 0.1 McF). The medium was analyzed for ADAM8 or ADAM10 activity using a FRET based assay, and the slope of the activity over time was calculated. Quantitative data are shown as mean + SD of three independent experiments. Statistical analysis was performed with one- sample t test (A-E) and one-way Anova (H-I). Asterisks represent statistical differences (* p < 0.05, ** p < 0.01, *** p < 0.001, **** p < 0.0001, n.s. not significant).

### Porphyromonas gingivalis induced ADAM8 activation correlates with tissue destructive activity

One characteristic of periodontal disease development is the overactivation of the immune response, in which neutrophils dominate the activate lesion, bridging both the innate and acquired immune systems (4). Infection of primary human neutrophils with *P. gingivalis* resulted in the release of exosomes carrying both the pro-form and remnant form of ADAM8 (Fig. 3A). In contrast, neutrophils showed a constitutive release of ADAM10 in exosomes under these culture conditions, which was not increased upon infection (Fig. 3B). Key factors for periodontal connective tissue destruction are MMPs such as MMP9, which have been shown to be regulated by ADAM8 (Cook et al., 2022). Zymography of patient samples revealed an increase in gelatinase-activity at higher disease stages (Suppl. Fig. 3A), which was confirmed by MMP9 ELISA measurements (Fig. 3C). Infection of primary human neutrophils revealed an increase in the cell-associated MMP9 expression (Fig. 3D), while the release content was dropping (Fig. 3E). However, in both cases MMP9 expression was significantly inhibited by BK-1361, and no effect by ADAM10 inhibition was observed. In contrast to neutrophils, MMP9 expression or release by keratinocytes was not affected by infection and inhibition, respectively (Suppl. Fig. 3B, C). Thus, neutrophil ADAM8 may regulate the tissue destructive activity in periodontal disease.

**Figure 3.**
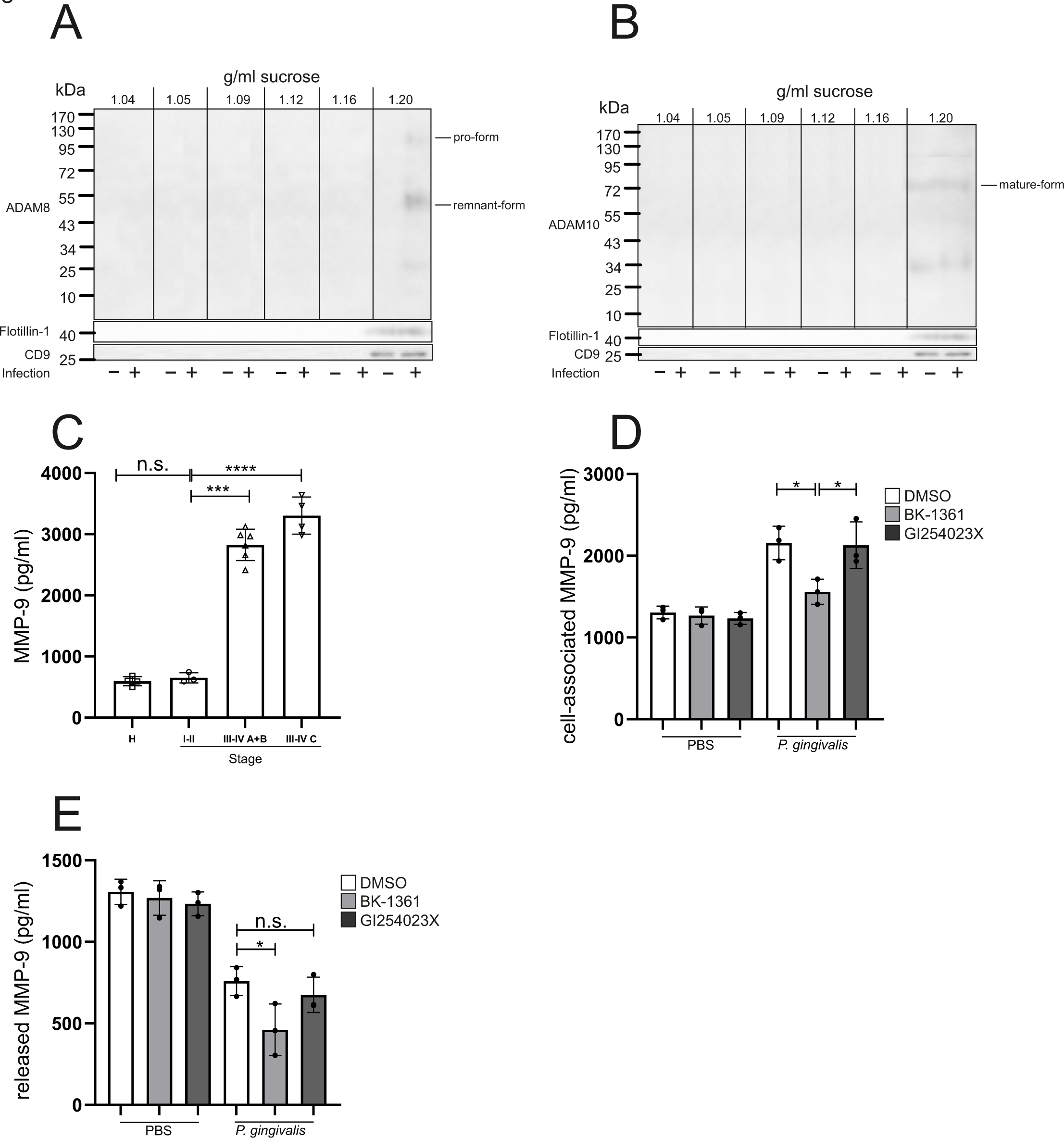
*P. gingivalis* induced ADAM8 activation correlates with tissue destruction. (A-B) Primary human neutrophils were infected with *P. gingivalis* (2 h, 0.1 McF) or left uninfected. The medium of the infected and non-infected cells was subjected to differential centrifugation and density fractionation for exosome purification. The resulting pellets of different density were analyzed by Western blot for the expression of ADAM8 and ADAM10 as well as Flottilin-1 and CD9 as exosomal markers. Representative images of three independent experiments are shown. (C) *S*ulcus fluid from patients with different disease stage was analyzed for MMP-9 expression by ELISA. (D-E) Primary human neutrophils were pre-incubated with 10 µM BK-1361, 10 µM GI254023X or 0.01% DMSO (vehicle control) for 30 min and infected with *P. gingivalis* (2 h, McF). Subsequently, the cells were lysed, and the cell lysate and the medium were analyzed for MMP-9 expression and release by ELISA. Neutrophil data are shown as mean +/- SD of three independent experiments. Statistical analysis was performed using one-way ANOVA followed by Tukey post hoc test. Asterisks without line represent statistically significant differences to healthy patients, asterisks with line between groups (* p < 0.05, *** p < 0.001, **** p < 0.0001, n.s. not significant).

### Cytokines may orchestrate the ADAM response in periodontal disease

Pathogens in the oral cavity lead to release of cytokines regulating not only the host immune response but also damage of the soft and hard tissue and osteocatalytic activity (3). The cytokine pool consists of the IL-1 and TNF family released by periodontal cells, myeloid cells and lymphocytes. To mimic this, we have chosen a combined stimulation with TNFα, IL-1β and IFNγ. After 24 h, keratinocytes showed an increase of ADAM8 pro-form, mature form and remnant form (Fig. 4A-C). Interestingly, only the pro-form remained elevated whereas the other forms returned to unstimulated levels upon long-term stimulation (48 h). For ADAM10, we observed a similar increase in protein expression, however not reaching significance due to high standard deviations (Fig. 4D,E). In contrast to ADAM8, the pro-form of ADAM10 was reduced after 48 h of stimulation. Analysis of the released exosomes revealed that 48 h stimulation led to an increase of the number of exosomes, indicated by an increase of the flotillin-1 and CD9 intensity (Fig. 5A, B). This was accompanied by weak expression of ADAM8 on these exosomes, but a strong increase of mature ADAM10 and a potential cleavage fragment (50 kDa). Activity measurements of ADAM8 and ADAM10 released from human primary keratinocytes and human neutrophils indicated an increase of ADAM8, which was inhibited by BK-1361. The increase of ADAM10 activity was inhibited by GI254023X in both keratinocytes and neutrophils. However, ADAM8 inhibition affected the ADAM10 activity in neutrophils only (Fig. 5 C-F).

**Figure 4.**
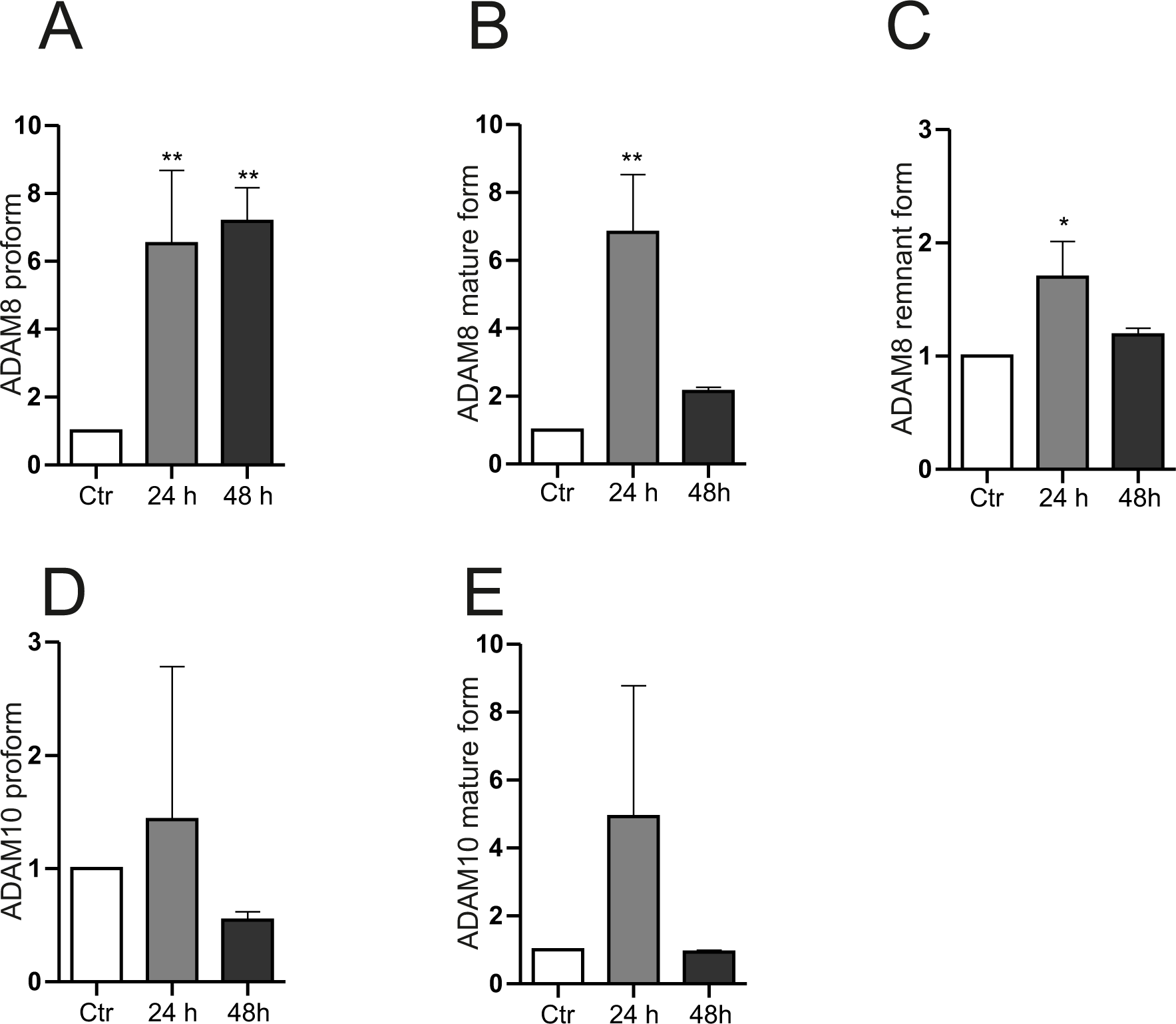
Cytokines regulate ADAM8 expression in human oral keratinocytes. (A-E) K2 cells were grown until confluency and stimulated with a combination of TNFα, INF-γ, and IL-1β (10 µM each) or sterile water (vehicle control) for 24 h and 48 h. Subsequently, the cells were lysed, and the samples were analyzed by Western blot using antibodies against the C-terminus of ADAM8 and ADAM10, respectively, followed by β-actin as loading control. Band intensity was quantified by densitometry and normalized to unstimulated cells. Data are shown as mean + SD of three independent experiments. Statistical analysis was performed by one sample t-test. Asterisks represent statistical differences (* p < 0.05, ** p < 0.01).

**Figure 5.**
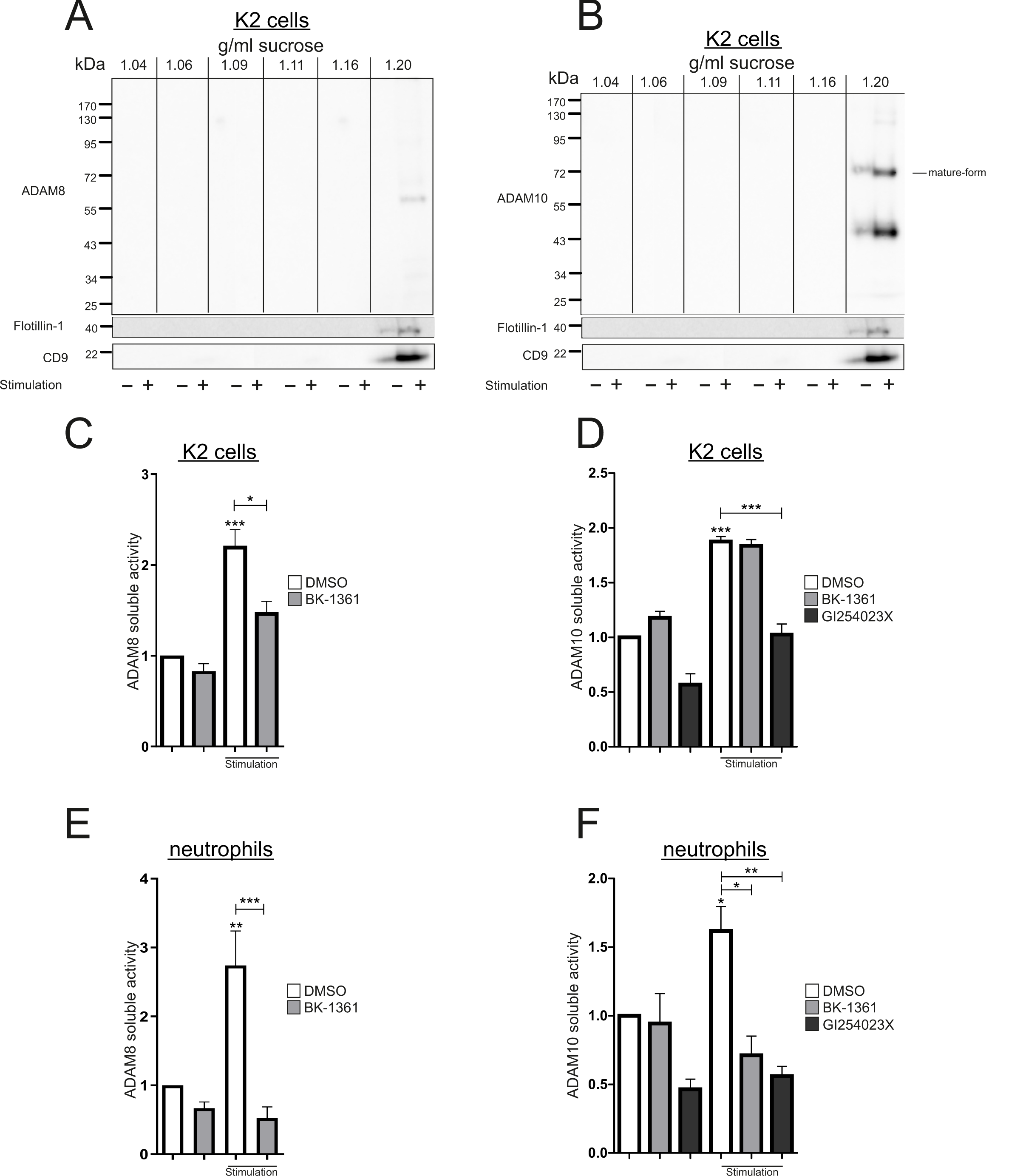
Regulation of extracellular release of ADAM8 and ADAM10 by cytokines. (A-B) K2 cells were grown until confluency and stimulated with a combination of TNFα, INF-γ, and IL-1β (10 µM each) or sterile water (vehicle control) for 48h. The medium of the stimulated and non-stimulated cells was subjected to differential centrifugation and density fractionation for exosomes purification. The resulting pellets of different density were analyzed by Western blot for the expression of ADAM8 and ADAM10 as well as Flottilin-1 and CD9 as exosomal markers. Representative blots of three independent experiments are shown. (C-F) K2 cells (C, D) or primary human neutrophils (E, F) were pre-incubated with 10 µM BK-1361, 10 µM GI254023X or 0.01% DMSO (vehicle control) for 30 min followed by stimulation with a combination of TNFα, INF-γ, and IL-1β (10 µM each) or sterile water (vehicle control) for 48 h. The medium was analyzed for ADAM8 (C, E) or ADAM10 (D,F) activity using a FRET based activity assay. Data are shown as mean + SD of three independent experiments. Statistical analysis was performed using one-sample t-test. Asterisks without line represent statistically significant differences to healthy patients, asterisks with line between groups (* p < 0.05, ** p < 0.01, *** p < 0.001).

Thus, bacterial infection and induced cytokine release stimulate a cell-specific release of ADAM8, orchestrating the tissue disrupting activity and resulting in novel proteolytic effector molecules such as ADAM10.

## Discussion

In the present study, we observed an increase of ADAM8 protein expression and activity in the sulcus fluid correlating with the periodontal disease stage. Both *P. gingivalis* infection and the resulting cytokine release may orchestrate the release of soluble ADAM8 by keratinocytes and neutrophils as soluble ectodomain or on exosomes, respectively. Furthermore, ADAM8 regulated the release of ADAM10 and MMP9, thereby potentially influencing wound healing and tissue destruction.

In previous studies, only the total ADAM8 protein expression was addressed (10, 12), which was similarly increased in our patient cohort and correlating with the disease stage. However, ADAMs are strongly regulated on the posttranscriptional level, resulting in activation and the generation of different protein forms. Indeed, we observed an increase of ADAM8 maturation as well as proteolytic activity, again correlating with the disease stage. It has been recently reported that ADAM8 exceeds proteolytic activity on ADAM10 and ADAM17 generating soluble ectodomains (9). Following these lines we observed a decrease of the membrane-associated forms of ADAM10 protein expression in the sulcus fluid. *In vitro*, we could confirm in both oral keratinocytes and neutrophils that ADAM8 activation is responsible for the release of ADAM10 proteolytic activity.

The observed soluble activity may result from the release of ADAM carrying EVs (17, 18) or the release of soluble ectodomains, which still exert proteolytic activity. ADAM8 requires trimerization resulting in the autocatalytic removal of the inactivating pro-domain and the release of its soluble ectodomain (19). This trimerization is inhibited by the cyclic peptide inhibitor BK-1361 (16). We observed an inhibition of ADAM8 activity by BK-1361 in cytokine stimulated oral keratinocytes only, whereas the inhibition of activity was weaker in infected cells. This was further reflected on the membrane-associated release of the mature form in EVs, which was absent upon infection only showing the remnant form. Furthermore, ADAM8 inhibition did not or only slightly affect the released ADAM10 activity. In contrast, neutrophil derived ADAM8 activity was inhibited by BK-1361 which further inhibited the ADAM8 release. Thus, keratinocytes seems to release ADAM8 as soluble ectodomain, whereas neutrophils are the cellular source of ADAM8 carrying exosomes.

Periodontal pathogens have developed strategies to evade the host immune defense (5). It was recently shown that Adam8-deficient mice are protected against bacterial lung infection, accompanied by a higher clearance capacity (20). Thus, an increase of ADAM8 protein expression in periodontal disease could be associated with less bacterial clearance and increase of bacterial burden. Furthermore, it has been reported that ADAM8 is important for MMP-9 release in the inflammatory tumor environment (17). MMP-9 is released during infection with *P. gingivalis* which is important for the breakdown of inflamed human pulp tissue (21), and it was shown that MMP inhibition protects against alveolar bone loss (22). In our setting, neutrophils could be identified as major source of the MMP9-release, which was inhibited by ADAM8. However, ADAM8 may not only modulate the tissue destruction and progression through MMP9 but also through release of ADAM10. It was shown that the soluble form of ADAM10 is capable to cleave fibronectin (9), with fibronectin fragments being associated with the periodontal disease status (23). Besides the release of soluble variants, we observed a decrease of the membrane-associated expression with increasing disease stage, which was also seen in our cell culture experiments. Cell-associated ADAM10 is required for the cleavage of growth factors such as epidermal growth factor ensuring proper wound healing (24) and contributes to the adhesion and migration of keratinocytes (25). Thus, the dysregulation of the cell-associated and extracellular ADAM proteolytic activity mainly driven by ADAM8 may be an essential regulatory element in periodontal disease onset or progression. Especially ADAM8, which seems to play subordinate role under physiological conditions and is mostly upregulated under pathophysiological conditions (16, 17, 20, 26, 27), offers a huge potential for local treatment and should be addressed in future translational studies.

## Materials and Methods

### Patient study

The study was approved by the ethics committee of the Landesärztekammer des Saarlandes (345/20 by DY and MH) and performed in accordance to the Declaration of Helsinki. Written informed consent was obtained from all individuals visiting either the Clinic for Dental Preservation, Periodontology and Preventive Dentistry of Saarland University Hospital or the dental office Reinstädtler and Hektor samples were pseudonymized during transfer from the clinics to the research area. Exclusion criteria were clinical signs of acute bacterial or viral infections, pregnancy, antibiotics within the last three month, use of mouth rinse, systemic diseases (e.g. rheumatoid arthritis, HIV), addictive disorders, low compliance, and missing consent. Healthy patients are those who routinely visited the clinics without sign of periodontal disease.

**Table 1:** Baseline characteristics of included patients.

| Demographic data | healthy patients (n =) | periodontitis patients (n =) |
| --- | --- | --- |
| age, male | 40 ± 19 | 51 ± 21 |
| age, female | 40 ± 22 | 50 ± 20 |
| gender, male, n (%) | 6 (40%) | 32 (60,4%) |
| gender, female, n(%) | 9 (60%) | 21 (39,6%) |
| stage I/II, grad A/B, male |  |  |
| stage III/IV, grad A/B, male | - | 7 (46,7%) |
| Stage III/IV, grad C, male | - | 10 (50%) |
| stage I/II, grad A/B, female | - | 3 (16,7%) |
| stage III/IV, grad A/B, female | - | 8 (53,3%) |
| Stage III/IV, grad C, female | - | 10 (50%) |

Patients were classified into stages (stage I-IV) and grades (grade A-C) according to the new classification of periodontitis from the EuroPerio9-Kongress which were introduced in 2018. The stage represents the severity of the disease and the grade shows the potential level of risk for disease progression. The severity is defined by Clinical Attachment Loss (CAL), radiographic bone loss and tooth loss. Complexity factors such as maximum depth of probing, bone resorption pattern, tooth mobility, furcation involvement and tooth loss. However, these can only worsen the staging and not change it to a less severe stage. Grading is done by determining bone loss in percent per age, whether and how much biofilm is present, longitudinal data on radiographic bone loss or CAL over a 5-year period, and other risk factors such as smoking and underlying diseases such as diabetes mellitus. We grouped the patients in stage I-II grade A+B, stage III-IV grade A+B und stage III-IV grade C.

### Sample preparation

Before sampling, the paper tips were weighed using precision scales and four paper tips were presented per reaction vessel. The surface of the selected teeth was cleaned with absorbent cotton rolls to remove debridement and saliva was removed by air blast. Relative drying was achieved in the maxilla by inserting an absorbent cotton roll vestibularly and in the mandible by absorbent cotton rolls vestibularly and lingually. The paper tips were inserted into the gingival pockets and left there for 30 s. The same procedure was followed for up to three further sampling sites of the same tooth and all tips were transferred together into a reaction vessel, so that the four previously weighed tips were always in the reaction vessel again. The samples were stored at 4°C for a maximum of seven days until further processing. By weighing the paper tips again, the volume of Gingiva-Crevicular-Fluid (GCF) was determined. The paper tips were trimmed so that the complete portion, which had become saturated with GCF, was in the reaction vessel. The remainder was discarded. For protein expression analysis by SDS-page and Western blot, the trimmed tips in the sample tube were eluted with ADAM lysis buffer (suppl. Table 2) as described below. For the activity assay, the tips were overlaid with 200µl of activity buffer (suppl. Table 2) as described below and incubated on ice for 20 minutes with repeated swirling for elution. Subsequently, the sample mixture was centrifuged for 10 minutes at 4°C and 4000xg and the supernatant was aliquoted into new sample vials and stored at -80°C.

### Antibodies, cytokines and inhibitors

**Table 2.** Cytokines, antibodies and inhibitors used in the present study.

| reagent | Company | Working concentration |
| --- | --- | --- |
| Recombinant Human TNF-α | Peprtech (Rocky Hill, NJ, USA) | 10 μM |
| Recombinant Human IFN-γ | Peprtech (Rocky Hill, NJ, USA) | 10 μM |
| Recombinant Human IL-1β | Sigma-Aldrich (Taufkirchen, Germany) | 10 μM |
| Goat polyclonal anti-human ADAM8 (Ectodomain) | R&D System, Wiesbaden, Germany | 0.2 μg/mL |
| rabbit polyclonal anti-human ADAM10 (C-terminus) | Invitrogen (Frankfurt, Germany) | 0.1 μg/mL |
| rabbit polyclonal anti-human GAPDH | Santa Cruz Biotech (Dallas, TX, USA) | 0.4 μg/mL |
| mouse monoclonal anti-human CD9 (MM2/57) | Invitrogen (Frankfurt, Germany) | 1 μg/mL |
| mouse monoclonal anti-Flotillin-1 | BD Biosciences (Heidelberg, Germany) | 0.25 μg/mL |
| Peroxidase-conjugated anti-mouse IgG | GE Healthcare (Chicago, IL, USA) | 1:20,000 |
| Peroxidase-conjugated anti-rabbit IgG | GE Healthcare (Chicago, IL, USA) | 1:40,000 |
| Peroxidase-conjugated anti-Gout IgG | GE Healthcare (Chicago, IL, USA) | 1:20,000 |
| BK-1361 | Peptide 2.0 (Chantilly, VA, USA) | 1 μg/ml |
| GI254023X | Merck Millipore (Darmstadt, Germany) | 10 μM |

**Table 2:** Buffers and solutions.

| Buffers and solutions | Composition |
| --- | --- |
| Lysis buffer | 20 mM TrisHCl<br>150 mM NaCl<br>5 mM EDTA<br>30 mM Na<sub>3</sub>Vo<sub>4</sub><br>5 mM DTT<br>1 mM PMSF<br>10 mM pNPP<br>1 mM benzamidine<br>10 mM glycerophosphate<br>1 mM Na<sub>3</sub>PO<sub>4</sub><br>1% Triton X-100 |
| ADAM8 FRET substrate activity buffer | 50 mM Tris<br>150 mM NaCl<br>10 mM CaCl<sub>2</sub><br>0.00006 % Brij<br>pH 8 |
| ADAM10 FRET substrate activity buffer | 25 mM Tris<br>10 mM CaCl<sub>2</sub><br>0.00006% Brij<br>pH 8 |
| SDS-sample buffer | 0.5 M Tris (PH: 6.8) |
|  | 45% Glycerol<br>5% SDS<br>0,25% Bromophenol blue<br>12.5% B-mercaptoethanol |
| Zymography washing buffer | 2.5% Triton X-100<br>50 mM Tris HCl<br>5 mM CaCl<sub>2</sub><br>1 µM ZnCl<sub>2</sub><br>H<sub>2</sub>O |
| Zymography incubation buffer | 1% Triton X-100<br>50 mM Tris-HCl<br>5 mM CaCl<sub>2</sub><br>1 µM ZnCl<sub>2</sub><br>H<sub>2</sub>O PH: 7.5 |
| Zymography staining solution | Methanol 40 %<br>Acetic acid 10 %<br>H<sub>2</sub>O 50 %<br>Coomassie Blue 0.5 % |
| Zymography destaining solution | Methanol 40 %<br>Acetic acid 10 %<br>H<sub>2</sub>O 50 % |

### Anaerobic bacteria preparation

The *P. gingivalis* strain DSM 20709 (syn. ATCC 33277) isolated from a human gingival sulcus was obtained from the German Collection of Microorganisms and Cell Cultures GmbH (DSMZ) and used throughout this study. DSM 20709 stocks were kept frozen at -80 °C. The bacteria were streaked out on Columbia agar plates supplemented with 5% defibrinated sheep blood and incubated under anaerobic conditions (85% N_2_, 10% CO_2_, 5% H_2_) at 37 °C for two weeks. A single colony was used to start a liquid culture in Brain Heart Infusion Broth (BHIB) and kept under anaerobic conditions at 37 °C for five days. Subsequently, the bacteria were centrifuged at 4,000xg for 5 min, resuspended in sterile PBS and adjusted to a McFarland of 1. The bacteria were used for *in vitro* infection experiments at 0.1 Mcfarland (McF).

### Cell culture and cell stimulation

Human keratinocytes (K2; gift from Dr. Miosge, University Göttingen) were cultured in Keratinocyte Growth Medium 2 (Promocell, Heidelberg, Germany) at 37°C 5% CO_2_ and 95% humidity following the manufacturer protocol. The cells were grown in 6-well plate or 15 cm dish (for exosome preparation) until confluence for 48-72 h. Blood from healthy donors were citrated and loaded slowly on Pancoll Human (PAN-Biotech GmbH, Aidenbach, Germany) for density fractionation. Subsequently, the layer containing erythrocytes and neutrophils was incubated with erythrocyte lysis buffer for 5-10 min at room temperature (RT). The neutrophils were cultured in RPMI1640 (+ 0.2% fetal calf serum (FCS)). For the inhibitor treatment, the cells were washed with PBS and pre-incubated for 30 min with ADAM8 cyclic peptide inhibitor BK-1361 or GI254023X, respectively, followed by *P. gingivalis* infection or cytokine stimulation as indicated in each figure legend.

### Western blot analysis

K2 cells or human neutrophils were lysed with 150-200 µl of lysis buffer (supp. Table 2) supplemented with 1X complete inhibitor (Roche) and incubated for 10 min at 4 °C. Subsequently, the lysates were centrifuged for 15 min at 16,000 g and the supernatant containing the proteins were subjected for protein determination using a commercial Bicinchoninic acid assay (BCA) kit (Thermo Fisher, Karlsruhe, Germany). 50 g of proteins were heat desaturated in SDS-sample buffer (supp. Table 2) and separated according to the molecular weight by SDS-PAGE as described in (8). The separated proteins were blotted onto a nitrocellulose membrane (Amersham Protran Premium 0.45 NC, GE Healthcare Life Sciences, Freiburg, Germany), blocked for the unspecific binding with 5% non-fatty milk and incubated overnight at 4 °C with the respective primary antibody. After washing, the membrane was incubated for 1 h with the secondary antibody, and the chemiluminescence signals were detected after adding the chemiluminescence substrate (PerkinElmer, Waltham, MA, USA) by luminescent image analyzer LAS3000 (Fujifilm, Tokyo, Japan) as described in (8). The intensity of the protein bands was quantified by selecting the same surface area in each band (after background correction) using AIDA Image Analysis software 4.27.039 (Elysia-raytest, Germany).

### Activity measurement

To assess the activity of the soluble ADAM8 in the patient samples as well as the release from K2 cells and human neutrophils, FRET-based substrates containing specific substrate cleavage sites were used.

K2 cells or human neutrophils were infected with *P. gingivalis* (0.1 McF) for 2 h or with combined cytokines (10 µM, TNF-α, IF- γ, IL-1β) for 24 and 48 h. the supernatant was centrifuged at 16,000 g for 10 min. Subsequently, 50 µl of the supernatant were transferred to a 96-well plate, followed by the addition of 50 µl of ADAM8 FRET substrate (Biozym Scientific GmbH, Hessisch Oldendorf, Germany) in FRET activity buffer(supp. Table 2) or 50 µl of ADAM10 FRET substrate. To determine ADAM8 or ADAM10 activity, the absorption was measured at 405 nm at 37°C every 2.5 minutes for 240 minutes using a Genios fluorescence reader (Tecan, Grödig, Austria). Data were calculated as the slop of the activity relative to the control.

### Gelatin zymography

Equal protein amounts isolated from sulcus fluid was loaded onto an SDS- polyacrylamide (5%) gel supplemented with 0.1% gelatin (from porcine skin, Sigma-Aldrich, Taufkirchen, Germany). Subsequently, the gel was washed with zymography washing buffer (supp. Table 2) twice, incubated with zymography incubation buffer (supp. Table 2); 10 min, 37 °C, with agitation), stained with zymography staining solution (supp. Table 2; 1 h, RT) and finally destained with zymography destaining solution (supp. Table 2; 10 min, RT). Areas of enzyme activity appear as white bands against a dark blue background.

### ELISA

Cell lysates, cell culture supernatants or sulcus fluid from patients with different disease stage were evaluated for MMP-9 content using ELISA Duo Set (DY911, R&D System, Wiesbaden, Germany.) following the manufactureŕs protocol.

### Exosome preparation

Isolate of exosomes from K2 cells or human neutrophils was performed as described in details in (8). Briefly, a total of 2 × 10^7^ cells were subjected to the indicated treatments. The conditioned medium was collected and subjected to a series of centrifugation steps: 10 min at 300xg, followed by 20 min at 1000xg, and 30 min at 10,000xg. After each centrifugation step, the resulting pellet was lysed in SDS buffer, while the supernatant was used for the next centrifugation step. The final supernatant was filtered through a 0.22 μm membrane filter, and the extracellular vesicles (EVs) were pelleted by centrifugation at 100,000xg for 1 h at 4°C using a Beckman rotor Type Ti50.2. The resulting EVs were washed with ice-cold PBS, pelleted again at 100,000xg for 1 h and dissolved in 100 µl and layered on a continuous gradient composed of layers of 2, 1.3, 1.16, 0.8, 0.5, and 0.25 M sucrose. After centrifugation at 100,000xg for 16 hat 4°C (Beckman Ti50.2 rotor), the density of each fraction was measured, and 1 ml of each fraction was pelleted again at 150,000xg for 4 h at 4°C using a Beckman TLA-55 rotor. The resulted sediment of each layer was then dissolved in SDS-sample buffer and analyzed by Western blot.

### Statistical analysis

Quantitative data are shown as mean+/- SD. Statistical analysis was performed using GraphPad PRISM 9.0 (GraphPad Software, La Jollla, CA). n-numbers and used statistical tests are given in the figure legends. A p-value < 0.05 was regarded significant.

## Acknowledgment

We thank Nina Schnellbach and Maria Rieseweber for technical support, and Jörg-Walter Bartsch and Nicolai Miosge for provision with BK-1361 and K2 cells, respectively. We thank the employees of the Clinic for Dental Preservation, Periodontology and Preventive Dentistry of Saarland University Hospital as well as those of the dental office Reinstädtler and Hektor for their support in acquiring the samples. We would also like to acknowledge the German Academic Exchange Service (DAAD) for their financial support.

## Author contribution

Hakon Heinze: Contributed to investigations, data analysis and writing the original draft

Federico Guillermo Gharzia: Contributed to investigations, data analysis and writing the original draft

Ahmad Aljohmani: Contributed to investigations, data analysis and writing the original draft. Ahmed Mohamed Mostafa Abdrabou: contributed to investigations.

Bashar Reda: Contributed to samples collection.

Sören Becker: Contributed to methodology, reviewed and edited the manuscript. Markus Bischoff: Contributed to methodology, reviewed and edited the manuscript.

Matthias Hannig: Contributed to conceptualization of the study, methodology, reviewed and edited the manuscript.

Daniela Yildiz: Contributed to conceptualization of the study, data analysis, methodology, reviewed and edited the manuscript.

## Abbreviations

ADAM, a disintegrin and metalloproteinase; BHIB, Brain Heart Infusion Broth; EVs, extracellular vesicles; ELISA, Enzyme-linked immunosorbent assay; FCS, Fetal calf serum; FRET, Fluorescence resonance energy transfer; GAPDH, Glyceraldehyde 3-phosphate dehydrogenase; IFN-γ, Interferon gamma, IL-1β, Interleukin 1 beta; MMP-9, matrix metalloproteinase 9; McF, Macfarland; *P. gingivalis, Porphyromonas gingivalis*; RT, Room temperature; TNF-α, Tumor Necrosis Factor alpha

---

**Supplementary Figure 1. Regulation of ADAM8, ADAM10 and ADAM17 in sulcus fluid of different disease stages.** Sulcus fluid from patients with different disease stage was investigated by Western blot to analyze the expression and maturation of ADAM8, ADAM10 and ADAM17 using antibodies against the C-terminal domains. GAPDH was used as loading control. Band intensity was quantified by densitometry. Representative blots are shown in A (ADAM8), C (ADAM10), and G(ADAM17). Quantitative data are shown as mean +/- SD. Statistical analysis was performed using one-way ANOVA followed by a Tukey post hoc test. (n.s. not significant).

**Supplementary Figure 2. *P. gingivalis* induced regulation of ADAM8 and ADAM10 expression in human keratinocytes**. (A-B) K2 cells were grown until confluency and infected with *P. gingivalis* (2 h, 0.1 McF). Subsequently, the cells were lysed, and the samples were analyzed by Western blot using antibodies against the C-terminal domains of ADAM8 and ADAM10. GAPDH was used as loading control. Representative blots of three independent experiments are shown.

**Supplementary Figure 3. Activity and presence of MMP9 in sulcus fluid and keratinocytes.** (A) Sulcus fluid samples from patients with different disease stages were subjected to a zymography. (B-C) K2 cells were grown until confluency, pre-incubated with 10 µM BK-1361, 10 µM GI254023X or 0.01% DMSO (vehicle control) for 30 min and infected with *P. gingivalis* (2 h, 0.1 McF). Subsequently, the cells were lysed and the cell lysate and the medium were analyzed for MMP-9 expression and release by ELISA. Data are shown as representative zymography in (A) and mean +/- SD of three independent experiments (B,C). Statistical analysis was performed using one-way ANOVA followed by a Tukey post hoc test. (n.s. not significant).

