## Supplementary figures and images for "Extracellular release of a disintegrin and metalloproteinases orchestrates periodontal disease severity"

### Supplementary figure 1

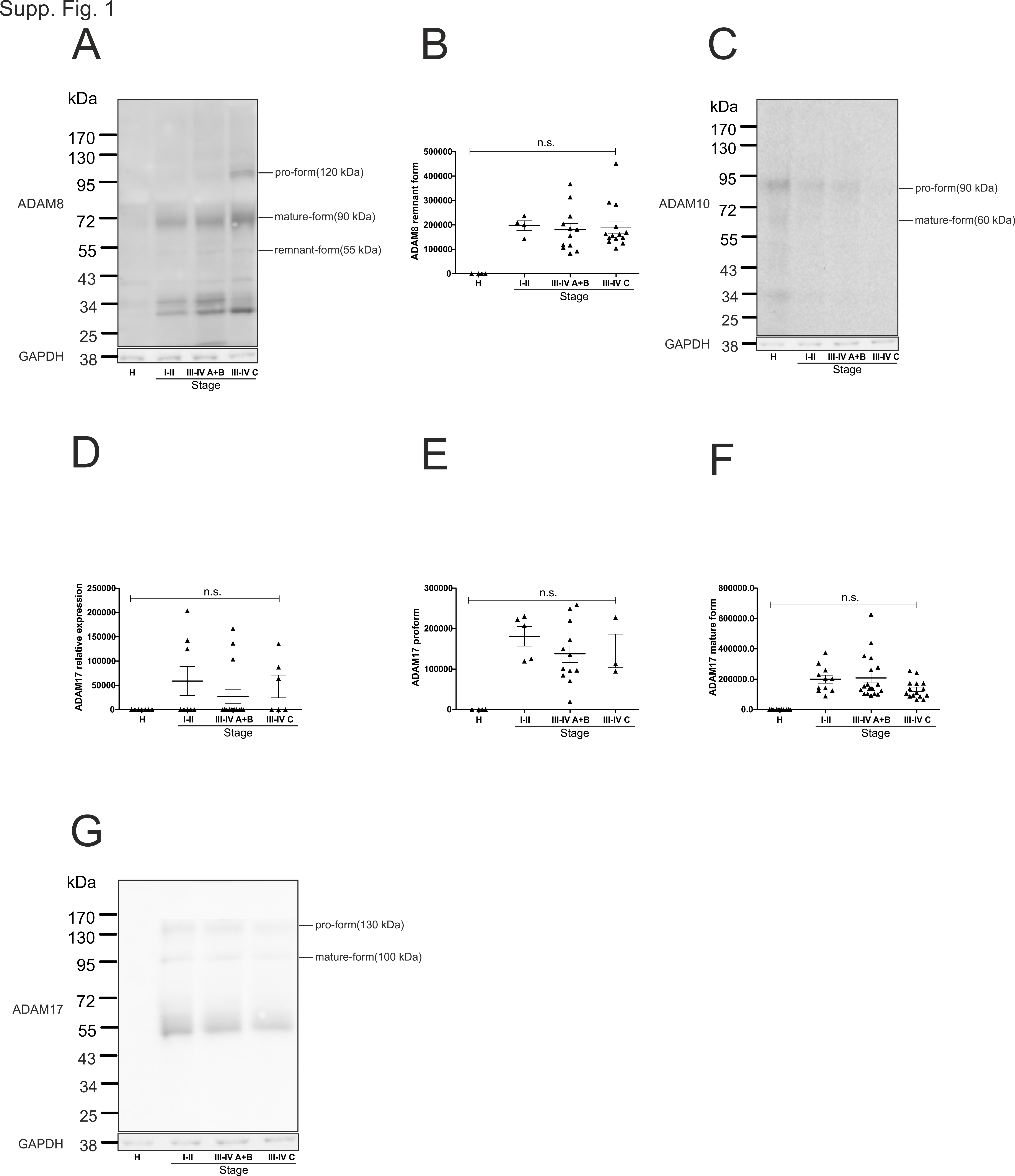

### Supplementary figure 2

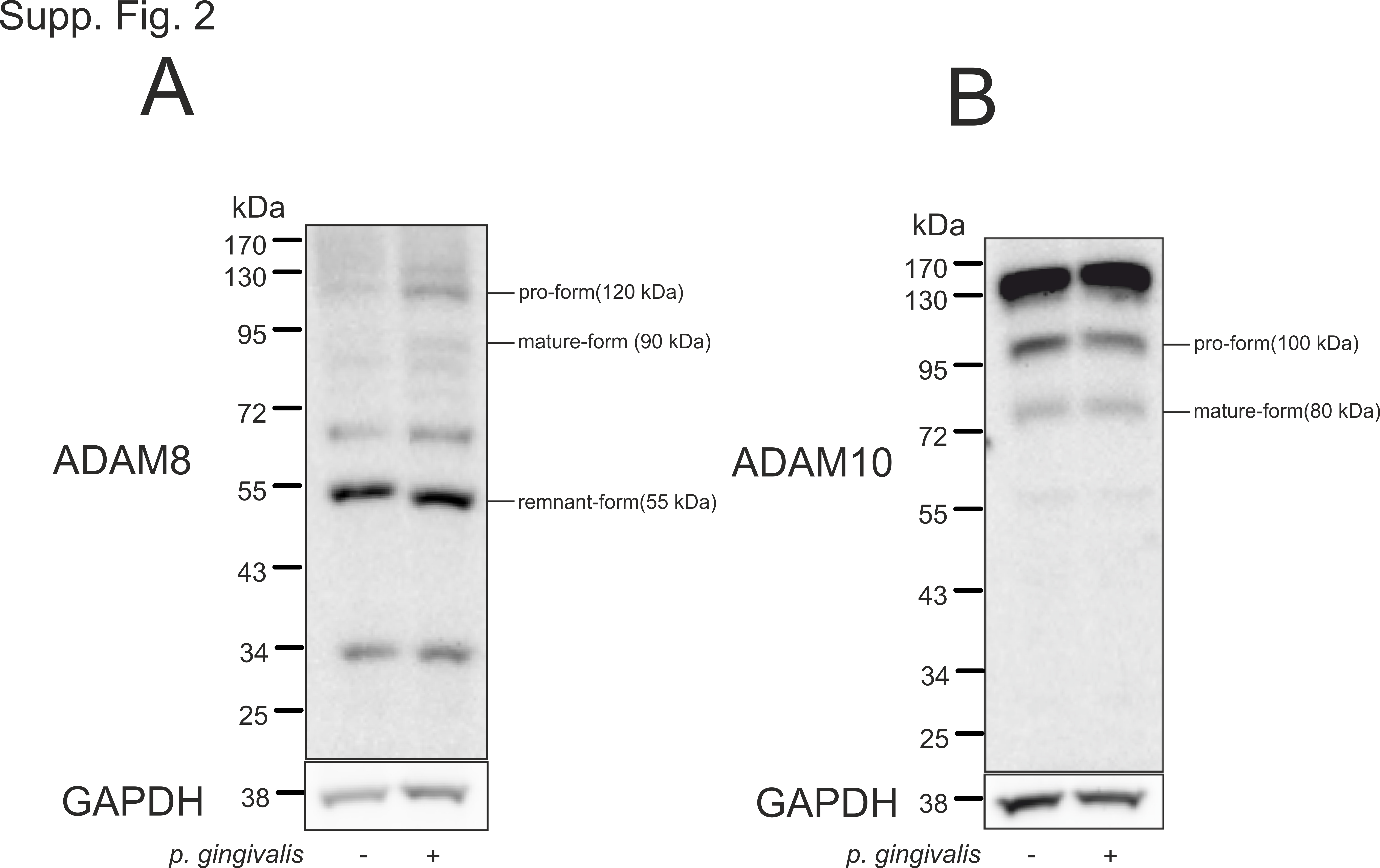

### Supplementary figure 3

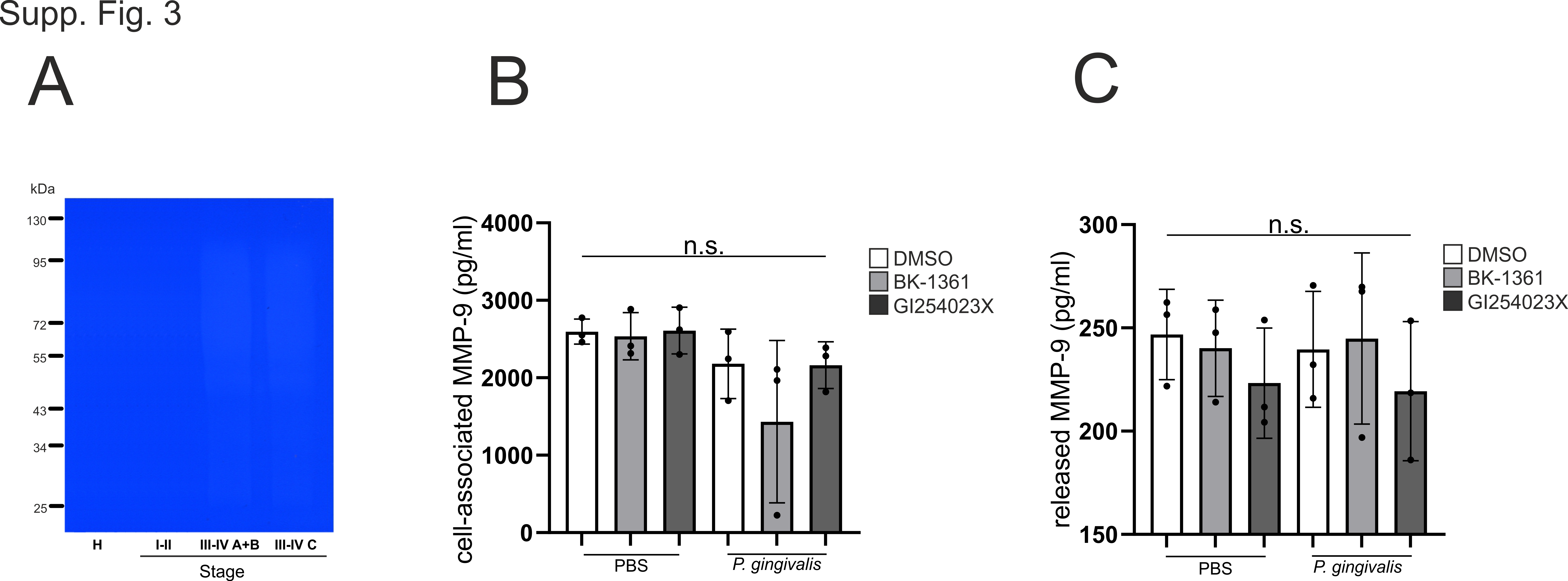
